# Accounting for animal movement improves vaccination strategies against wildlife disease in heterogeneous landscapes

**DOI:** 10.1101/2021.10.15.464547

**Authors:** Katherine M. McClure, Guillaume Bastille-Rousseau, Amy J. Davis, Carolyn A. Stengel, Kathleen Nelson, Richard B. Chipman, George Wittemyer, Zaid Abdo, Amy T. Gilbert, Kim M. Pepin

## Abstract

Oral baiting is used to deliver vaccines to wildlife to prevent, control, and eliminate infectious diseases. A central challenge is how to spatially distribute baits to maximize encounters by target animal populations, particularly in urban and suburban areas where wildlife like raccoons (*Procyon lotor*) are abundant and baits are delivered along roads. Methods from movement ecology that quantify movement and habitat selection could help to optimize baiting strategies by more effectively targeting wildlife populations across space. We developed a spatially explicit, individual-based model of raccoon movement and oral rabies vaccine seroconversion to examine whether and when baiting strategies that match raccoon movement patterns perform better than currently employed baiting strategies in an oral rabies vaccination zone in greater Burlington, Vermont, USA. Habitat selection patterns estimated from locally radio-collared raccoons were used to parameterize movement simulations. We then used our simulations to estimate raccoon population rabies seroprevalence under currently used baiting strategies (actual baiting) relative to habitat selection-based baiting strategies (habitat baiting). We conducted simulations on the Burlington landscape and artificial landscapes that varied in heterogeneity relative to Burlington in the proportion and patch size of preferred habitats. We found that the benefits of habitat baiting strongly depended on the magnitude and variability of raccoon habitat selection and the degree of landscape heterogeneity within the baiting area. Habitat baiting improved seroprevalence over actual baiting for raccoons characterized as habitat specialists but not for raccoons that displayed weak habitat selection similar to radio-collared individuals—except when baits were delivered off roads where preferred habitat coverage and complexity was more pronounced. In contrast, in artificial landscapes with either more strongly juxtaposed favored habitats and/or higher proportions of favored habitats, habitat baiting performed better than actual baiting, even when raccoons displayed weak habitat preferences and where baiting was constrained to roads. Our results suggest that habitat selection-based baiting could increase raccoon population seroprevalence in urban-suburban areas, where practical, given the heterogeneity and availability of preferred habitat types in those areas. Our novel simulation approach provides a flexible framework to test alternative baiting strategies in multiclass landscapes to optimize bait distribution strategies.

## INTRODUCTION

Oral baiting is commonly used in invasive species and infectious disease management to deliver toxicants, biologics or pharmaceuticals to wildlife across landscape-level scales (Savarie et al. 2001, Howald et al. 2007, Slate et al. 2009). For instance, vaccine-laden baits targeting disease reservoir populations are typically deployed at large spatial scales as part of disease management efforts to control or eliminate pathogens like rabies virus, bovine tuberculosis, and sylvatic plague (Mencher et al. 2004, Müller et al. 2015, Nugent et al. 2016). Vaccine baiting has also been used both experimentally and operationally to protect endangered animal populations threatened by infectious disease, including Allegheny woodrats at risk from raccoon roundworm and Ethiopian wolf populations threatened by rabies virus (Haydon et al. 2006, Smyser et al. 2013, Sillero-Zubiri et al. 2016). In invasive species management, baited lethal toxicants and fertility control vaccines have been deployed over large spatial scales to reduce vertebrate pest populations like brushtail possums (Tompkins and Ramsey 2007). Across diverse management objectives, a central challenge in wildlife baiting is how to deliver and distribute baits to maximize bait encounter and consumption by free-ranging target animal populations, particularly across heterogeneous, multiclass landscapes.

Bait consumption by wildlife requires placing baits in areas where animals will find and consume them within hours to days. One strategy to improve encounter rates (i.e. rates of exposure to baits by wildlife, given bait availability) is to draw from methods in movement ecology to examine how target populations use the landscape and incorporate animal-habitat association patterns into the spatial design of baiting programs (Beasley et al. 2015). This baiting strategy could benefit from recent advancements in wildlife tracking and biotelemetry technologies that have led to a significant increase in the availability of wildlife movement data, along with the concurrent development of analytical frameworks to quantify animal movement and habitat selection across gradients of land use and habitat complexity (Nathan et al. 2008, Tomkiewicz et al. 2010, Long and Nelson 2013, Kays et al. 2015). For instance, habitat selection analyses are a set of statistical approaches that link individuals to their environment to identify habitat selection patterns, including avoidance and preferential use of a resource or habitat type. One common approach uses resource selection functions (RSF; Boyce et al. 2002) to estimate habitat selection using land cover or environmental conditions coupled with GPS location data, and can be used to predict the relative probability of occurrence of animals on the landscape (Johnson 1980). These methods could help to improve wildlife baiting effectiveness by incorporating movement behavior and space use patterns into stratified baiting designs to increase bait encounter and uptake rates among target populations. There is a growing appreciation that habitat selection analyses can be used to improve bait encounter rates by better understanding where animals are most likely to optimally forage (e.g. Berentsen et al. 2013, Schneider et al. 2019). Yet habitat selection patterns have been rarely explicitly included in invasive species and disease management and planning (Boyer et al. 2011, Beasley et al. 2015).

Raccoon rabies virus variant (RRV) is a wildlife zoonosis in North America that has been managed using oral vaccine baiting (Slate et al. 2020). Rabies virus (RV) is a single-stranded RNA virus in the genus Lyssavirus that is transmitted between animals by direct contact, and can lead to a central nervous system infection that is invariably fatal across mammals (Rupprecht et al. 2002, Wunner 2007). In North America, at least eight distinct rabies variants are maintained in wild carnivore reservoir species, including RRV (Gilbert 2018), which accounts for the majority of rabies virus exposures to humans and spillover infections among animals in the United States (Wallace et al. 2014, Pieracci et al. 2019). Along the eastern U.S. seaboard, RRV is enzootic in raccoons (*Procyon lotor*), an urban-adapted meso-carnivore that is ubiquitous in both rural and developed landscapes (Hadidian et al. 2010). Coordinated RRV management in the U.S., led by the U.S. Department of Agriculture Animal and Plant Health Inspection Service National Rabies Management Program (USDA-APHIS NRMP), utilizes oral rabies vaccination (ORV) as the primary strategy to reduce the susceptible fraction of individuals in target raccoon populations and contain the westward spread of RRV from the eastern U.S. (Slate et al. 2009). Oral rabies vaccines used in the U.S. are enclosed in plastic sachets or blister packs coated in fishmeal or sweet vanilla attractant that induce immunity when ingested by raccoons and other target wildlife (Slate et al. 2009, Moore et al. 2017, Blanton et al. 2018, Gilbert et al. 2018b). Sustained annual ORV led to the successful elimination of canine RV variant from coyotes in the U.S. and red fox RV variant throughout most of the European Union (Sidwa et al. 2005, Muller and Freuling 2018). Achieving sufficient population-level vaccination coverage in North American raccoon populations has been challenging, however, particularly in urban areas where raccoons reach high densities and bait delivery is typically accomplished by hand baiting from vehicles (Riley et al. 1998, Gilbert and Chipman 2020).

Components of baiting strategies that can be manipulated to improve bait consumption within budgetary, personnel, and logistical constraints include bait density (baits/km^2^) and the spatial arrangement of baits applied to the landscape (where baits are placed, e.g., along roads or targeting specific land cover classes) (WHO 2018, Gilbert and Chipman 2020). The objective of this study was to investigate conditions of raccoon habitat selection in heterogeneous landscapes that could increase consumption of oral rabies vaccine baits and population-level antibody seroconversion. Specifically, we evaluated raccoon habitat selection and avoidance conditions which could improve the effectiveness of ORV baiting strategies in urban-suburban landscapes. To address these questions, we first characterized raccoon habitat preferences using resource selection functions estimated from GPS location data from raccoons captured within an ORV zone in Burlington, Vermont, USA. Next, we developed a spatially explicit, individual-based model (IBM) of raccoon movement and bait consumption to predict population-level rabies antibody seroconversion, or seroprevalence, under different spatial baiting designs and habitat selection patterns (e.g. data-based versus more specialist or generalist). We applied our simulation framework to an 81 km^2^ gridded landscape in Burlington as well as artificial landscapes that varied in landscape composition (i.e. the identity and proportion of land cover types) and patch size of landcover types. We hypothesized that using data-based movement and habitat selection patterns of raccoons to inform the spatial distribution of baits would increase vaccine bait encounters and seroconversion for raccoons relative to standard baiting conditions. Our novel simulation framework provides insight into how habitat selection behavior could guide wildlife or disease management strategies using oral baiting and could be readily extended to model diverse movement behaviors across a variety of heterogenous landscapes to determine optimal bait-distribution strategies.

## METHODS

### Study area and data collection

#### Study area

Our study area is located within a 222 km^2^ ORV management zone in the greater Burlington, Vermont area designed to control RRV in target populations of raccoons, striped skunk (*Mephitis mephitis*), gray and red foxes (*Urocyon cinereoargenteus* and *Vulpes vulpes*, respectively), bobcats (*Lynx rufus*), and coyotes (*Canis latrans*) (Fig. 1A). The area is characterized by suburban and urban land-use types embedded within a matrix composed of undeveloped land and open space, deciduous forests, and farmland. The meso-carnivore species composition in this area includes several species that can act as bait competitors like the domestic cat (*Felis catus*) and Virginia opossum (*Didelphis virginianus*) (Slate et al. 2020).

**Figure 1.**
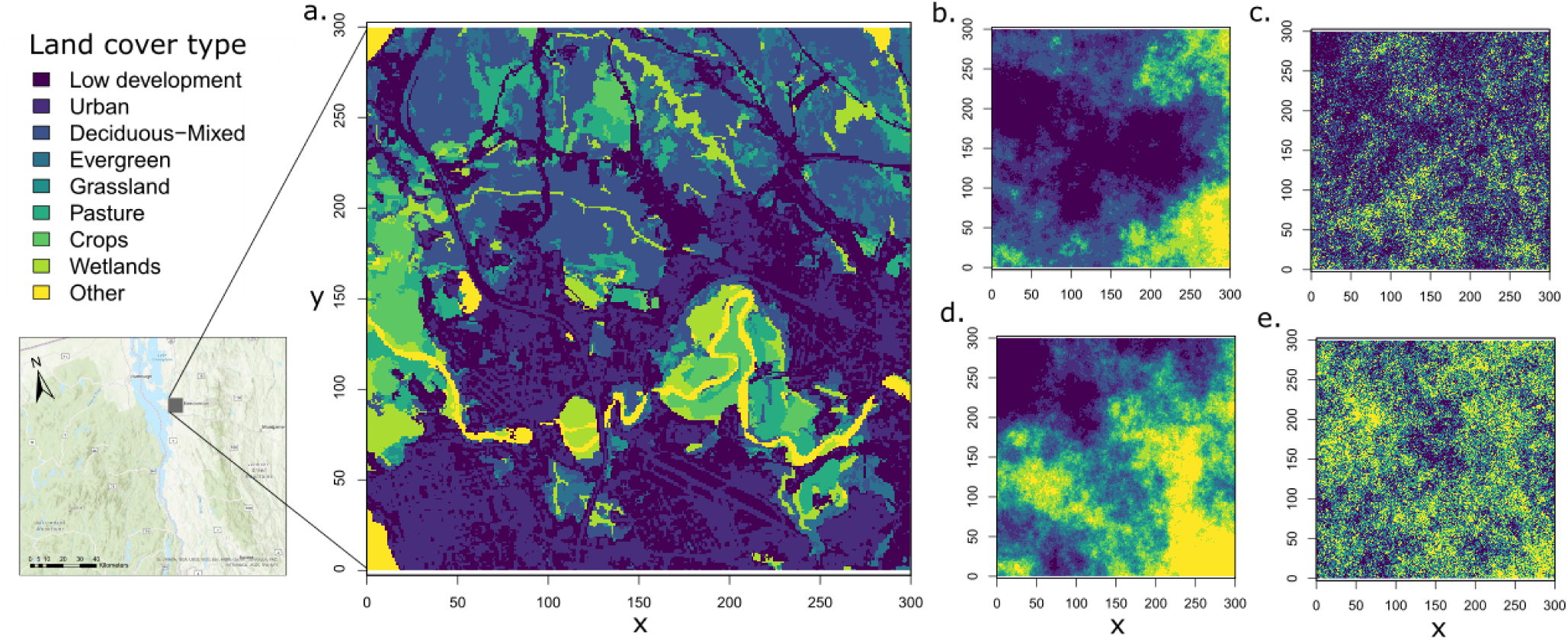
Actual and artificial landscapes used in movement and bait seroconversion simulations. a) The 81km^2^ study area within a current ORV zone in Burlington, Vermont, USA used in simulations (with regional area map shown, bottom left), comprising nine landcover types at a 30 m spatial resolution. In a separate analysis, four types of multiclass, artificial landscapes of similar area and spatial resolution were used in simulations, each consisting of two parameters: patch size (P) and landscape composition (C). Relative to Burlington, simulated landscapes had: b) similar patch sizes and amount of preferred habitat; c) smaller patch sizes; d) more preferred habitat; e) smaller patch sizes and more preferred habitat.

#### Actual ORV baiting

During 2016-2017, ORV bait distribution was planned and executed within the Burlington ORV zone by the NRMP. The ORV baits were distributed along roads and deployed by hand and vehicle within habitat defined operationally as baitable using the following procedure. The Burlington ORV zone was divided into six planning subunits, each averaging 37 km^2^. Each subunit was baited at a target density of 150 baits/km^2^ across landcover types characterized by the Multi-resolution Land Characteristics Consortium’s National Land Cover Data 2011 (NLCD; Homer et al. 2015). The number of baits required to meet density targets at the subunit level were calculated using a habitat-based algorithm, defined operationally as an off-time calculator, that was based on expert opinion of raccoon ecology by the ORV team at the USDA-APHIS NRMP. The algorithm reduces bait distribution in some land cover types while remaining land cover types are baited at a target density of 150 baits/km^2^ for the subunit. Specifically, no baits are placed in open water (NLCD land cover category = 11) or barren (NLCD = 31) land cover types. In low, medium, and high intensity development and pasture/hay land cover types (NLCD = 22 – 24, 81; respectively), baits are placed at lower target densities (97.5, 52.5, 15, and 75 baits/km^2^, respectively). During bait delivery operations in August 2016 and 2017, point location data were collected for each bait deployed across the Burlington ORV zone using Point of Interest GPS technology (“NRMP baiting” hereafter; Appendix S1: Fig. S1; Tsi GL-770, Transystem Inc., Hsinchu City, Chinese Taipei).

Our simulated focal landscape encompassed an 81 km^2^ gridded landscape within the larger Burlington ORV zone, and had a 900 m^2^ subunit resolution (30 m by 30 m grid cells; described in detail below). The average bait density distributed by NRMP using the bait delivery algorithm was 110 baits/km^2^ across the study area we defined, given the land cover composition within this area and the associated land cover-specific target densities (see Appendix S1: Table S1 for the composition and total area of NLCD land cover types within our study area used in bait delivery planning). To reflect NRMP average baiting levels in our defined study area, we implemented an average bait density of 110 baits/km^2^ for baiting simulations. Thus the average bait densities for NRMP-based and habitat selection-based baiting were similar but the densities within particular landcover types were different. This approach allowed for comparison between NRMP conditions and our simulated habitat selection-based baiting strategy, which by design lacked most land cover-based restrictions except for those related to habitat selection patterns (details below). We used a finer spatial scale in our simulations than used for operational bait delivery because we estimated habitat selection at this smaller spatial scale and because it enabled the simplifying assumption that raccoons and baits located within the same grid cell equated to a raccoon encountering a bait.

#### Raccoon capture and tracking

Raccoons were captured prior to fall ORV during 2016 using cage traps (Tomahawk model 608, Tomahawk Live Trap, Wisconsin, USA). We fitted 25 individuals with Q4000E GPS collars (Telemetry Solutions, California, USA) equipped with very high frequency (VHF) beacons. During July through September, individual raccoon GPS relocation data, or fixes, were collected every 30 minutes to 6 hours. This study followed guidelines outlined by the Institutional Animal Care and Use Committees of the University of New Hampshire (#160901) and USDA-APHIS National Wildlife Research Center (QA2669). In this area, population-level virus neutralizing antibody seroprevalence was estimated through pre- and post-ORV serum sampling of target meso-carnivore populations, as described in Gilbert et al. (2018b).

### Simulation model

#### Overview

The workflow of our analyses and simulation framework is described in Fig. 2. First, we analyzed raccoon GPS data (Fig. 2.1A) in order to: 1) calculate distances moved by individual raccoons at a 30-minute time step (i.e., step lengths; the distribution of movement distances at each time step for each individual) and 2) estimate resource selection coefficients for each raccoon using landcover data from the 30 m resolution raster NLCD land cover data (Homer et al. 2015; Fig. 2.1B and C). We then developed an IBM of raccoon movement and bait encounter informed by these analyses that matched the spatial resolution of the habitat selection analysis and the temporal resolution of the GPS data collection (Fig. 2.2D). To investigate the effects of raccoon habitat selection, we considered a continuum of selection strength ranging from more general to more specialized, which were both relative to the estimated habitat selection of radio-tracked raccoons, and modeled individual- and population-level habitat selection, in separate simulations. We held individual-level movement parameters constant across different habitat selection conditions to focus on the effect of magnitude and variation in habitat selection on predicted bait encounter and seroconversion (Fig. 2.2D).

**Figure 2.**
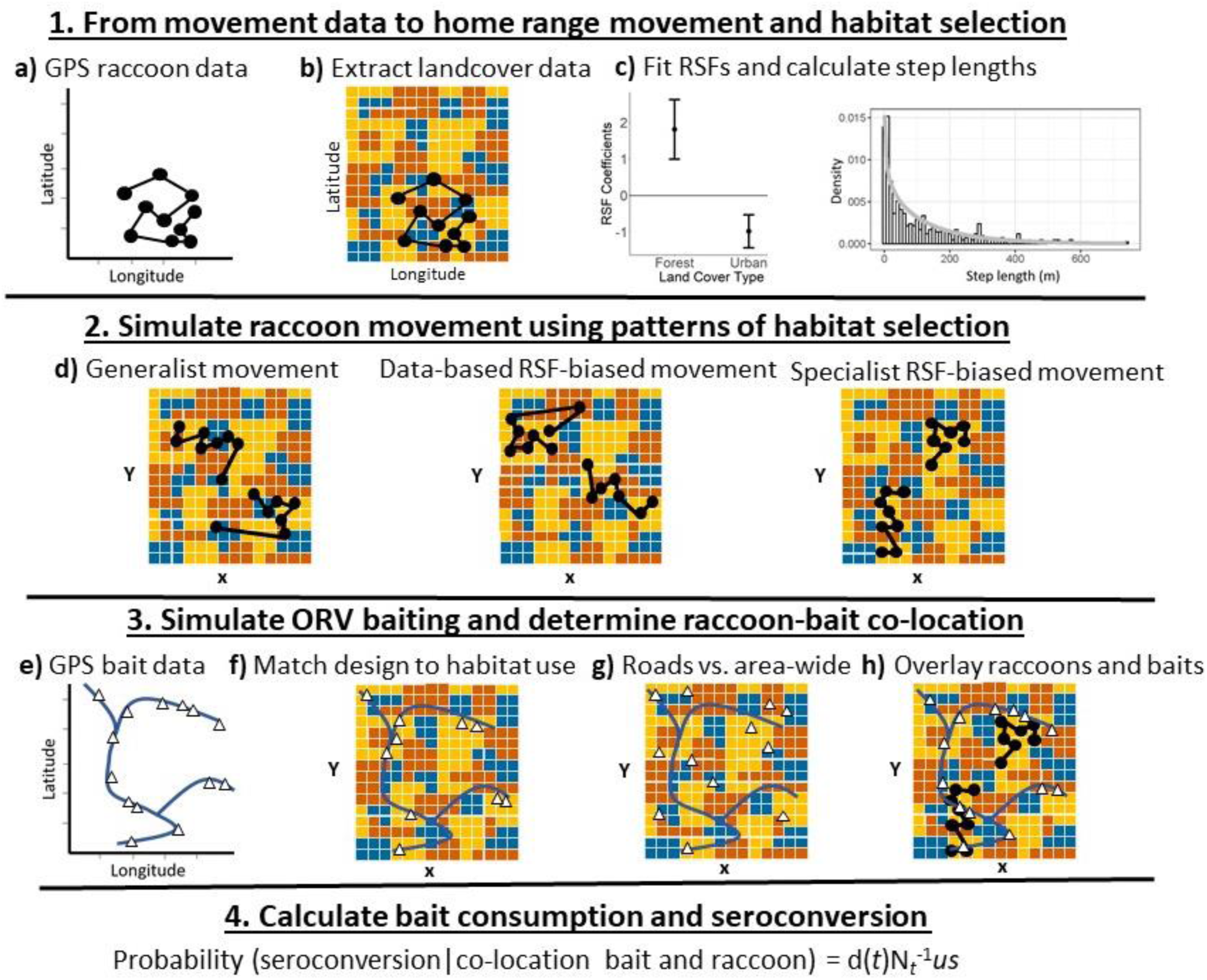
Simulation workflow. To simulate raccoon movement and oral-baited rabies vaccine seroconversion, we first collected a) GPS relocation data from 25 wild-caught raccoons captured within an oral rabies vaccination (ORV) vaccination zone in Burlington, Vermont, USA during July - September, 2016. We used b) 30 m resolution raster land cover data reclassified into nine land cover types to c) fit resource selection functions (RSF) to individual-level raccoon relocation data and fit a gamma distribution to individual-level 30-minute interval distances moved (step lengths; in m). d) Next, we simulated raccoon movement across gridded landscapes composed of nine land cover types according to different assumptions of habitat selection behavior, each in separate simulations. Then we simulated different ORV baiting designs (e - g) and h) spatially overlaid raccoons and baits. We calculated the probability of seroconversion in individual raccoons given within-cell vaccine bait co-location using a discrete daily decay function (d*(t)*, see text), consumption rate given co-location (*u* = 0.5), seroconversion rate given consumption (*s* = 0.9), and randomly assigned baits for potential consumption if multiple raccoons were present in a cell (N = total within-cell raccoons).

Next, to explore the importance of raccoon habitat selection on bait encounter using different baiting designs, we simulated baiting designs that differed relative to NRMP baiting operations in three components of baiting designs: 1) the spatial arrangement of baits (i.e., the proportion of baits in different landcover types and road-limited versus area-wide baiting), 2) coverage (evenness of bait distribution across the study area), and 3) density (baits/km^2^) (Fig. 2.3E - G). We then overlaid simulated baiting designs on raccoon movement scenarios and estimated the proportion of raccoons that encountered and consumed a vaccine bait, resulting in seroconversion (rabies antibody seroprevalence of the simulated population; main output of the simulation model). Bait consumption and seroconversion involved three processes (Fig. 2): 1) co-location of a raccoon and bait within the same grid cell, 2) bait consumption given co-location that accounted for non-target and conspecific bait competition and reduced bait availability over time, and 3) seroconversion given bait consumption. We estimated seroprevalence for each set of conditions relative to operational baiting and data-based raccoon movement in the area (Fig. 2.3H and 2.4).

Finally, to investigate how landscape heterogeneity influences the effectiveness of baiting designs based on raccoon movement ecology, we compared outcomes conducted on the landscape in our study area of Burlington, Vermont to those from identically sized, artificial landscapes that differed in the proportion and patch size of landcover types preferred by radio-tracked raccoons in Burlington (Fig. 1). Simulations and data analysis were conducted in program R v3.6.2 (see Data S1 for simulation code; R Core Team 2019). We describe each component of the analysis in detail below.

#### Estimation of habitat selection and movement distances from GPS data

We used resource selection functions (RSF; Boyce et al. 2002) to evaluate raccoon third-order (within home range) habitat selection using a use-available design (Johnson, 1980). We compared GPS locations of 25 individual raccoons to available points (n = 10,000) generated within each raccoon’s seasonal late summer/autumn home range. We used 95% kernel density estimation (KDE) to estimate home range size during the three months of telemetry data collection. Because this 3 month duration may not represent stationary raccoon home ranges, we opted for a simple (and potentially more conservative) estimate of home range size that better reflects actual use instead of including probable future use in the delineation (e.g. autocorrelated kernel density estimation; Noonan et al. 2019). Landcover was extracted for all locations from NLCD land cover data (R package raster; Hijmans 2021). We reclassified land cover into nine categories that we assumed were functionally relevant to raccoon ecology as described in Davis et al. (2019): open space and low density development, medium to high density development, deciduous and mixed forest, coniferous forest, shrub and grassland, pasture, crops, wetlands, and other (Appendix S1: Table S1). Individual-level RSF coefficients for each land cover type relative to open space and low density development were estimated using an exponential regression (fitted using a logistic regression; McDonald 2013) using the IndRSA package (https://github.com/BastilleRousseau/IndRSA; Bastille-Rousseau and Wittemyer 2019). We estimated population-level RSF coefficients using generalized linear mixed models with a binomial distribution, a logit link, and individual as a random intercept (Gillies et al. 2006, package lme4 in R; Bates et al. 2015). We estimated distributions of step lengths—distances moved by individuals at a 30-minute time scale—by fitting a gamma distribution to the distances (continuous measure of distance in meters; with gamma distribution parameters shape and scale; Appendix 1: Table S2) between GPS relocation fixes of each tracked raccoon at a 30-minute interval.

#### Simulation of raccoon movement

We simulated discrete-space raccoon movement across 30 m spatial resolution gridded landscapes (30 m x 30 m; 900 m^2^ cell size, the same spatial resolution as the land cover data) composed of nine landcover types using a biased random walk algorithm (Uhlenbeck and Ornstein 1930). Raccoons moved among grid cells according to rules governed by three independently parameterized processes, each given equal weight, including raccoon density, home range attraction, and habitat selection informed by landcover and our habitat selection analyses (Fig. 2A - D). Movement was modeled at a 30-minute time step to match the temporal resolution (i.e. fix rate) of raccoon GPS data. For each raccoon at each time step, a continuous distance was randomly drawn from the step length distribution. For distances < 15 m, individuals stayed within the cell at time *t* + 1 whereas for step lengths > 15 m, a set of potential destination grid cells surrounding the individual’s current cell location were defined. The product of three processes was used to weight the cell-level probability of movement to each potential destination cell: 1) habitat selection, 2) home range attraction, and 3) conspecific avoidance relative to host density. The probability of selecting a cell based on habitat selection was weighted using RSF coefficients, calculated at the individual or population-level depending on the movement scenario, as:

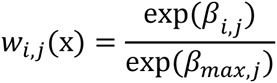

where w(*x*) is the relative habitat selection strength at cell *x* of individual or population *j* for landcover type *i* in geographical space, β*_i_* is the selection coefficient for landcover type *i*, and β*_max_* is the largest estimated habitat selection coefficient for individual or population *j*. Home range attraction, *h*, was modeled to constrain raccoon movement within a home range area, and was calculated as a discrete approximation of a negative exponential function, where the squared distance (in m) of cell *k* from the home range centroid was calculated as:

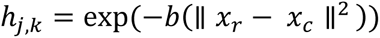

where *b* is the decay rate, *x* are points in Euclidean space, *c* is the centroid of cell *k* and *r* the home range center point of individual *j*. Finally, we modeled conspecific density-dependent effects on movement behavior, which we calculated as the inverse of the count of raccoons in a cell in the previous time-step. Simulations were performed for 1440 time-steps, equivalent to one month of movement.

To explore the effects of magnitude and variation in raccoon habitat selection on bait encounter across similar conditions of habitat availability, we simulated movements in the gridded landscape in greater Burlington that differed in the strength of individual- and population-level habitat selection (i.e., data-based, generalist, or specialist) (Table 1). Data-based raccoon movement reflected habitat selection patterns estimated using GPS data from radio-tracked raccoons in greater Burlington, which suggested that raccoons in this area exhibit substantial individual variation in habitat selection resulting in weak population-level habitat selection (Table 1 and Appendix S1: Table S2). Generalist movement reflected a theoretical case in which animals neither preferred or avoided any land cover type; it was thus equivalent to random movement with respect to land cover but was constrained by conspecific density and home range attraction. Specialist movement behaviors were also theoretical but informed by data in that they were modeled by doubling the data-based estimated RSF coefficients for the preferred landcover types for radio-tracked raccoons (preferred types included NLCD-classified deciduous (41) and mixed forest (43) land cover classes and wetland areas including woody wetlands (90) and herbaceous wetlands (95); Table 1 and Appendix 1: Table S2). In the individual-level data-based selection scenario, we incorporated estimated resource selection coefficients for individual radio-tagged raccoons (Appendix 1: Table S2), assigned randomly to each simulated individual at the start of each simulation and in tandem with an individual’s associated movement parameters. The population-level data-based scenario reflected the average habitat selection response of radio-tagged raccoons, drawn from the population-level habitat selection analysis, such that selection was constant among individuals (Appendix 1: Table S2). Under our population-based specialist scenario, all individuals exhibited constant, strong selection, while individuals in the individual-based specialist scenario exhibited variation in habitat selection strength among individuals (Appendix 1: Table S2). Comparing results across the continuum of habitat specialization and variability allows for inference about how baiting designs can be optimized based on knowledge of local animal movement behavior and the influence of individual-level movement heterogeneity (e.g. (McClure et al. 2020)). Each habitat selection scenario was applied at three raccoon densities that represent a realistic range of raccoon densities in suburban-urban landscapes (5, 15, 30 raccoons/km^2^; Urban 1970, Moore and Kennedy 1985, Slate et al. 2020). Each raccoon movement scenario (habitat selection scenario × raccoon density) was repeated 100 times.

**Table 1.** Raccoon habitat selection patterns and bait component scenarios implemented in simulations. High bait density = 110 baits/km^2^ and low bait density = 55 baits/km^2^.

| Raccoons |  |  | Baiting |  |  |  |
| --- | --- | --- | --- | --- | --- | --- |
| Initiation of raccoon on the landscape | Habitat selection | Density (inds/km<sup>2</sup>) | Distribution: landcover targeting | Distribution: road targeting | Coverage (% grid cells with baits) | Density (baits/km<sup>2</sup>) |
| Random | Generalist | 30, 15, 5 | Habitat-selection based vs NRMP off-time calculator | Road-limited (NRMP) vs area-wide | 3% (NRMP 2016) vs 6% (NRMP 2017) | High (NRMP) vs low |
| Based on similar population-level resource use | Data-based, population average | 30, 15, 5 | Habitat-selection based vs NRMP off-time calculator | Road-limited (NRMP) vs area-wide | 3% (NRMP 2016) vs 6% (NRMP 2017) | High (NRMP) vs low |
| Near favored habitat (deciduous and wetland) | Specialist, population average | 30, 15, 5 | Habitat-selection based vs NRMP off-time calculator | Road-limited (NRMP) vs area-wide | 3% (NRMP 2016) vs 6% (NRMP 2017) | High (NRMP) vs low |
| Based on similar individual-level habitat selection | Data-based, individual variation | 30, 15, 5 | Habitat-selection based vs NRMP off-time calculator | Road-limited (NRMP) vs area-wide | 3% (NRMP 2016) vs 6% (NRMP 2017) | High (NRMP) vs low |
| Near favored habitat (deciduous and wetland) | Specialist, individual variation | 30, 15, 5 | Habitat-selection based vs NRMP off-time calculator | Road-limited (NRMP) vs area-wide | 3% (NRMP 2016) vs 6% (NRMP 2017) | High (NRMP) vs low |

#### Initialization of raccoon movement simulations

Raccoon populations were initialized on the landscape for each simulation iteration using the habitat selection patterns described above. Initial locations were generated at random for generalist movement conditions (Appendix S1: Fig. S2B). For data-based scenarios, we evaluated the composition of each individual and population-based range in terms of landcover type (Appendix S1: Fig. S2A). We extracted the proportion of each landcover type around each grid cell within 20 cells or a 600 m radius (e.g. corresponding to the average estimated Burlington raccoon home range size, data not shown) and initiated individuals proportionally to the similarity (i.e. how similar the area is in terms of proportion of each landcover) between observed home range composition and surrounding grid cells. For specialist scenarios, initial locations occurred in areas with higher abundance of selected landcover types (woody and herbaceous wetland and deciduous or mixed forest) within a 600 m radius (Appendix S1: Fig. S2C).

#### Simulation of bait consumption and seroconversion

Baits were delivered within cells on day 20 of each movement simulation. We assumed each raccoon co-located with a bait in a grid cell had a bait consumption probability of 0.5 based on the expert opinion of NRMP (Fig. 2). Spatial overlap between baits and target animals on the landscape are necessary for bait uptake by target animals, but other factors such as bait competition both from conspecifics and non-target species also influence bait uptake probability (Elmore et al. 2017, Slate et al. 2020). We accounted for bait competition from conspecifics by randomly selecting one raccoon to encounter the bait when multiple raccoons were co-located in a cell, leading to a bait consumption probability of less than 0.5 when raccoon densities were high within grid cells. To account for competition from non-target carnivores like opossums (Slate et al. 2020) and decreasing bait palatability over time, we included a discrete decay function for bait availability in which 30% of baits within a grid cell were randomly removed every 12 hours (discrete equation: n_h =_ n_init_ *.70*^h^*^/12^, where *h* = hour, n = number of baits, init = initial time step; similar to bait disappearance rates reported by Smyser et al. (2013)), with a maximum bait availability time of 10 days. Lastly, we assumed that seroconversion probability, given consumption, was 0.9 to reflect immunogenicity following oral contact with the vaccine (Brown et al. 2012, Gilbert et al. 2018a). For each simulation, we extracted the proportion of raccoons that seroconverted, or rabies antibody seroprevalence.

#### Baiting designs

We modified three components of baiting strategies: spatial arrangement, coverage, and bait density. Spatial arrangement included two conditions for how the baits were placed relative to landcover data: 1) using NRMP bait distribution that incorporated the habitat-based off-time calculator (NRMP baiting), or 2) using raccoon habitat selection patterns as described above (described hereafter as habitat baiting). To simulate habitat baiting targeting specialist and data-based raccoons, baits were distributed proportionally to resource selection of a given landcover type according to population-level RSF coefficients as described above. For strict generalists that did not display habitat preferences or avoidances, baits were randomly distributed among all landcover types except open water. Across habitat-varying spatial delivery strategies, each condition was simulated by either constraining baiting delivery to roads only (road-limited baiting; actual condition) or by delivering baits over the entire study area (area-wide baiting; alternative condition). We examined two bait densities: one that was 110 baits/km^2^ across all landcover types including those with reduced or no baiting applied as described above (high bait density; equivalent to 150 baits/km^2^ in NRMP-defined baitable habitats as described above) and a lower density condition of 55 baits/km^2^ across all landcover types (low bait density; equivalent to 75 baits/km^2^ in NRMP-targeted habitats). Finally, we compared two levels of grid cell coverage, although this factor is not routinely measured for spatial planning of bait delivery by NRMP. Low coverage included 3% (1233/41100) of grid cells with baits (similar to NRMP baiting pattern in 2016) while high coverage included 6% (2466/41100) of grid cells with baits (similar to NRMP baiting pattern in 2017). We caution that operational baiting strategies are implemented at a larger spatial resolution than the scale at which we modeled bait delivery. However, because NRMP baiting varied among years in overall landscape coverage, we used the concept of grid cell coverage to examine the evenness of baiting effort across space in an ORV area. The full set of raccoon movement × bait strategy simulation scenarios, 240 in total, are described in Table 1.

#### Artificial landscapes

To explore how landscape characteristics and road structure in Burlington influenced our results, we conducted a separate set of simulations on artificial landscapes that varied in the proportion of land cover types preferred by raccoons (landscape composition) and the clustering of those land cover types across the study area (patch size based on fractal dimension). We generated neutral landscapes using fractional Brownian motion with the R package NLMR (Sciaini et al. 2018). We simulated four artificial landscapes with the same road structure and spatial extent as our study area that differed in one or both dimensions of patch size and landscape composition relative to Burlington. Specifically, these four artificial landscapes had: 1) similar patch sizes and proportion of all landcover types as Burlington (Fig. 2B), 2) smaller patch sizes of preferred habitat (i.e. more fragmented) but with the same landscape composition as observed in Burlington (Fig. 2C), 3) similar patch sizes as in Burlington, but with an increase in preferred habitat relative to Burlington (i.e. an equal proportion of each landcover type and thus higher proportion of preferred habitat; Fig. 2D), and 4) smaller patch sizes of preferred habitat and an increase in preferred habitat relative to Burlington Fig. 2E).

For each artificial landscape, we simulated raccoon movement with either data-based or specialist habitat selection patterns estimated from radio-tagged raccoons captured in Burlington, as described previously. We assumed similar individual-level habitat selection for movement simulations in these artificial landscapes (i.e. not the population average nor population-level) along with conspecific density and home range attraction movement constraints. We created a different landscape for each iteration and used NRMP conditions for raccoon and bait density parameters (15 raccoons/km^2^, high bait density application) because simulation results on the Burlington landscape did not indicate interactions in these parameters. For each habitat selection condition, we compared: 1) habitat baiting to NRMP baiting, 2) road-limited to area-wide baiting, and 3) coverage of 3% versus 6% of study grid cells (Table 1).

### Statistical analyses

We used a linear mixed effects model with a logit link to analyze the relative impacts of baiting strategy features on the primary model outcome: rabies antibody seroprevalence. All covariates were considered as categorical (i.e. “dummy” variables). Replicate was added as a random intercept because each scenario was repeated 100 times. Simulation parameters (habitat selection patterns by raccoons, raccoon density, bait distribution by landcover, bait density, bait coverage, and road-only versus area-wide baiting) were included as main effects. An interaction between raccoon habitat selection and bait distribution by landcover, and a series of three-way interactions of this interaction with all other parameters were also added. Heat maps were produced from the model predictions to illustrate the impact of all parameters and their interactive influences relative to NRMP conditions (data-based habitat selection by raccoons and NRMP-based baiting from 2016: high bait density, road-limited baiting, 3% coverage, which includes the NRMP’s off-time algorithm for determining distribution of baits to different landcover types). Simulations based on artificial landscapes were analyzed similarly. In this case, an interaction between habitat selection by raccoons and landscape type was added and a series of three-way interactions with all other parameters.

## RESULTS

### Habitat selection and step length patterns

At the population level, raccoons displayed selection for woody and herbaceous wetlands, deciduous and mixed forest, and grassland and shrub land cover types relative to developed open space and low intensity development. Raccoons exhibited substantial individual variation in resource selection with some individuals selecting resources avoided by other raccoons (Appendix 1: Table S2). Individual response was especially strong for deciduous and mixed forest with 56% (14/25) of individuals selecting it and 28% (7/25) of individuals avoiding it, and for wetlands with 52% (13/25) and 32% (8/25) of individuals selecting and avoiding it, respectively. Raccoons consistently avoided medium and high intensity development relative to open space and low intensity development (92%, 23/25). Individual variation was also present in step length (Appendix S1: Table S2).

### Effects of bait distribution

Our model was scaled to predict post-ORV population-level seroprevalence among raccoons at similar levels to that measured seasonally by NRMP in Burlington during 2016 and 2017 (44% [N = 195] and 27% [N = 234] respectively; our model: 35 [34 - 36 CI]% under 2016 baiting conditions; Fig. 3A). Contrary to our predictions, for data-based raccoon movement, seroprevalence predictions were similar or slightly lower under habitat-based baiting relative to NRMP baiting for road delivery (Fig. 3A, B, D), but with area-wide baiting the habitat-based baiting led to higher seroprevalence as we predicted (Fig. 3C). However, if raccoons were either habitat generalists or specialists, seroprevalence was substantially higher compared to NRMP baiting when baiting matched habitat selection patterns (up to 21% higher for specialist) under all conditions (Fig. 3). Overall, using population-level movement rules increased seroprevalence predictions for all combinations compared to individual-level movement rules (Appendix S1: Fig. S3), suggesting that higher individual-level variation in habitat selection decreases seroprevalence under any given baiting design.

**Figure 3.**
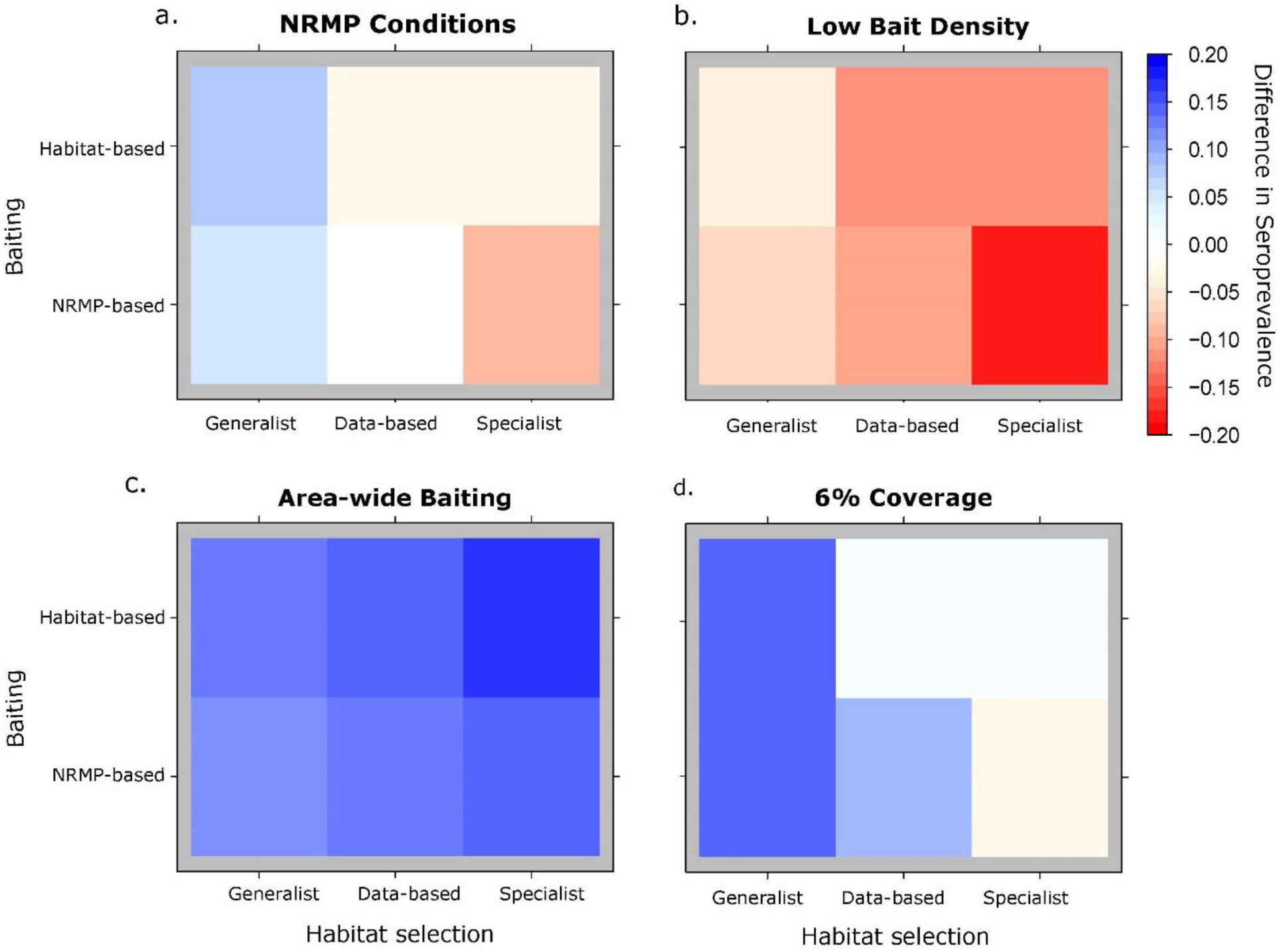
Effects of baiting design and raccoon habitat selection on predicted rabies seroprevalence. Plots show absolute difference in seroprevalence for different baiting designs relative to NRMP conditions (white block in a, with predicted absolute seroprevalence of 0.35). Red shades indicate that the design leads to lower seroprevalence relative to NRMP baiting conditions while blue shades indicate higher seroprevalence. Rows in each panel correspond to the spatial distribution of baiting strategies across land cover types (i.e. proportion of baits distributed to different landcover types). Habitat-based and NRMP-based baiting stratifies baiting using habitat selection patterns and an algorithm drawn from decades of raccoon ORV experience by the NRMP, respectively. Columns correspond to raccoon habitat selection behaviors. Labels on top of the panels indicate changes in the baiting strategy relative to NRMP conditions with a) representing standard NRMP delivery and distribution, b) lower bait density, c) bait baits were distributed across the full area rather than on roads, and d) spatial coverage was doubled, to 6%.

### Effects of bait density and coverage

Bait density and coverage affected overall seroprevalence, but not which baiting design was best relative to raccoon movement (Fig. 3B, D). Decreasing the bait density from our high to low treatment decreased seroprevalence by 11% under otherwise standard NRMP baiting conditions (Fig. 3B). Increasing grid cell baiting coverage from 3% to 6% increased seroprevalence by 8.5%. However, although predicted seroprevalence of the habitat-based strategy was similar to the NRMP baiting strategy under both bait density scenarios, it unexpectedly performed worse than the NRMP baiting strategy along roads when coverage was increased from 3% to 6%.

### Effects of raccoon density

Raccoon density had a strong impact on predicted seroprevalence but did not influence which strategy led to greater seroprevalence (Fig. 4), indicating that the baiting strategies acted similarly across the raccoon densities we examined (Appendix S1: Table S3). Under the observed baiting and movement conditions, seroprevalence was predicted to be 3% higher when raccoon density was decreased from 15 to 5 raccoons/km^2^ and 3% lower when raccoon density was doubled, from 15 to 30 raccoons/km^2^. This indicates that while raccoon density is a significant factor driving overall seroprevalence, it does not affect whether habitat-based or NRMP-based baiting is more effective.

**Figure 4.**
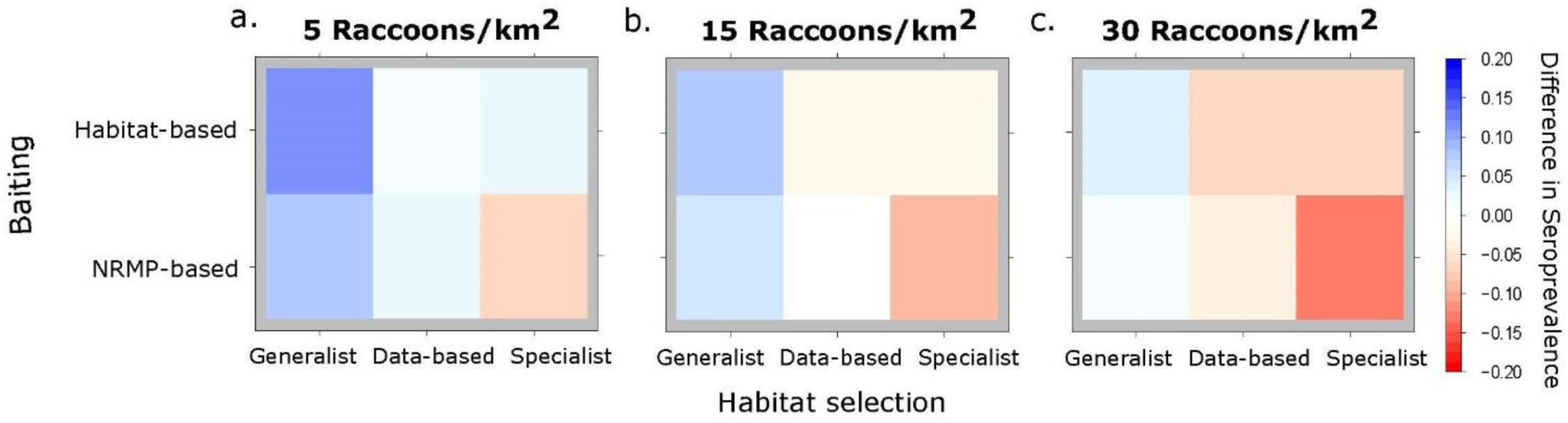
Effects of raccoon density. Panels are the same as in Fig. 3 except that NRMP conditions are in the lower middle of panel b (white block, predicted absolute seroprevalence of 0.35) and each panel reflects results for different raccoon densities: a) 5 individuals/km^2^, b) 15 individuals/km^2^, and c) 30 individuals/km^2^.

### Effects of landscape composition and structure

The result that habitat-based baiting was only advantageous when habitat selection was strong or with area-wide baiting suggested that some of our results may have been impacted by the specific road structure relative to landscape composition and patch size in Burlington. Visual inspection of the Burlington landscape revealed that most roads do not intersect raccoon-preferred habitat and the habitat roads do intersect is relatively homogenous (i.e. large patch size). Thus, we compared seroprevalence for the habitat-based and NRMP baiting strategies using data-based versus specialist raccoon habitat selection in artificial landscapes, while assuming the same area and road structure as in Burlington (Fig. 5). Here, we found that when there was a higher proportion of raccoon-preferred habitat or smaller patch sizes, habitat-based baiting was better, even when raccoons displayed weak selection for preferred land cover types (i.e. reflecting data-based habitat selection patterns) and using road-based delivery methods (Fig. 5A & B). We also found that the importance of area-wide baiting for increasing seroprevalence was reduced in these other landscapes (higher proportion of raccoon-preferred habitat, smaller patch size) relative to Burlington (where patch sizes were large and raccoon-preferred habitat was limited; Fig. 5C). In other words, in these artificial landscapes, the amount of benefit gained from habitat-based baiting depended on the composition and patch size of raccoon-preferred habitat relative to raccoon movement patterns, especially in proximity to roadways. Increasing coverage from 3% to 6% increased seroprevalence rates by up to 18% in all the artificial landscapes and by 5% in Burlington (Fig. 5D), suggesting that targeting habitat along roads more systematically may improve seroprevalence under many different landscape conditions.

**Figure 5.**
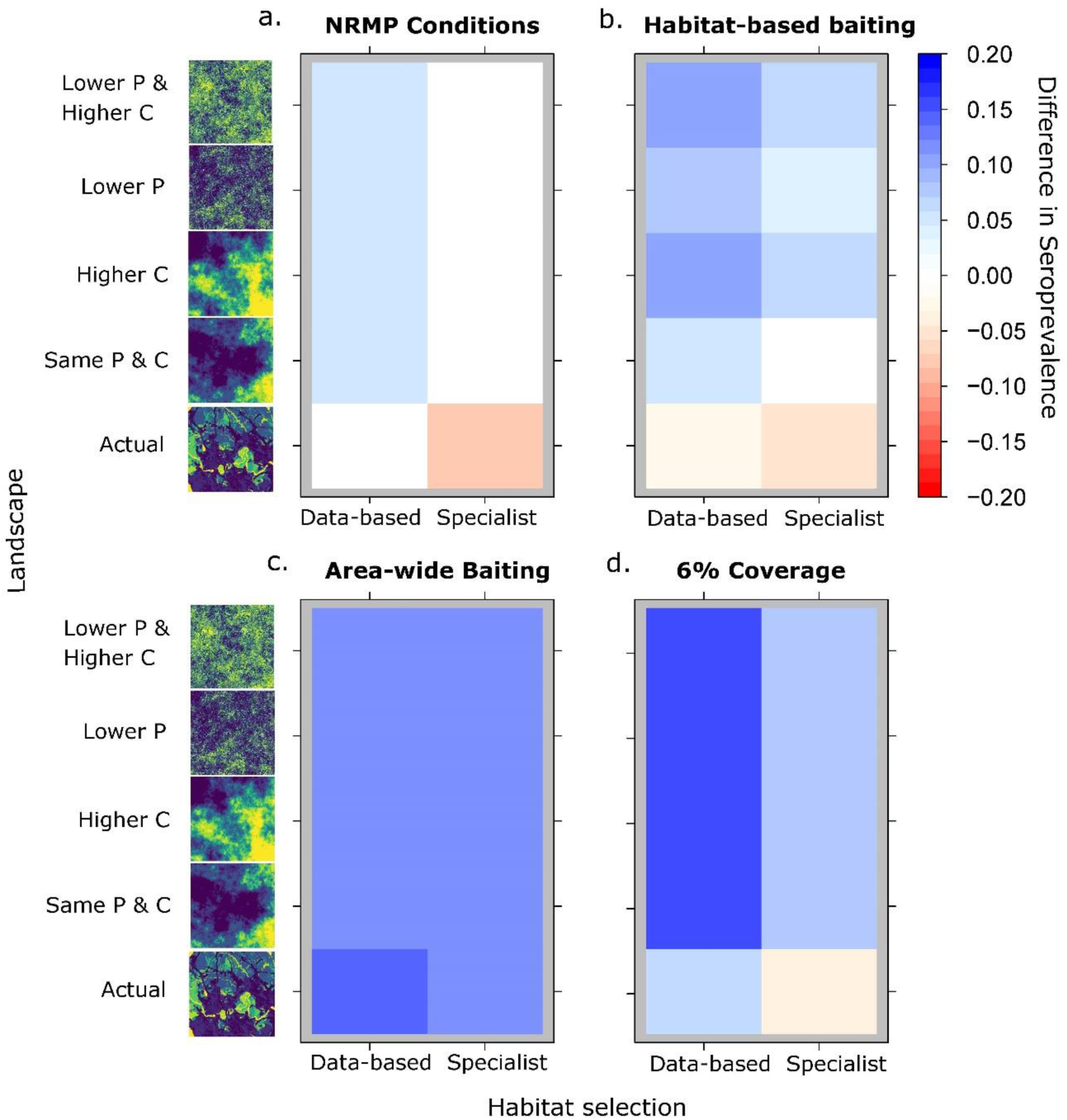
Effects of landscape composition and patch size. Rows within panels describe different landscape conditions: Burlington, Vermont, USA ( “actual”) and four simulated landscapes comprised of two landscape metrics relative to Burlington (described in Fig. 1): patch size (P) and proportion (C) of preferred habitat of raccoons. Columns correspond to raccoon habitat selection behavior (‘data-based’ describes weak habitat selection whereas ‘specialist’ describes strong habitat selection). Colors represent absolute differences in seroprevalence relative to the NRMP conditions (white square in lower left corner in a; 0.35 predicted seroprevalence) as described in Fig. 3. Labels on top of the panels indicate changes in the baiting strategy relative to NRMP conditions.

## DISCUSSION

The effectiveness of oral rabies vaccination programs targeting raccoons can be hindered by challenges associated with landscape-level bait delivery and processes that influence bait uptake rates by raccoons across heterogeneous landscapes. We developed a novel simulation framework that incorporates movement behavior of free-ranging raccoons, realistic multiclass landscapes, and spatial bait data from previous ORV efforts to explore if and how raccoon movement behavior could be integrated to increase oral vaccine baiting-associated population immunity to rabies virus. We identified raccoon habitat-specialization conditions and landscape characteristics that determine how different components of baiting strategies can be altered to optimize population seroconversion to ORV in urban and suburban areas. We found that habitat-based baiting increased seroprevalence when raccoons showed strong habitat selection patterns or when landscapes had substantial areas of raccoon-preferred habitat and were more heterogeneous (small patch sizes), suggesting habitat baiting using selection patterns could be beneficial in some but not all landscape contexts. Relatedly, we show that individual-level variability in raccoon habitat preference and avoidance patterns decreased seroprevalence using any baiting design. Finally, increasing bait coverage, bait density, or expanding the distribution of baits beyond roads led to higher seroprevalence under all baiting designs, but these components of baiting strategies did not affect whether or not it is advantageous to do habitat-based versus NRMP-based baiting.

In our models, habitat-based baiting in the greater Burlington ORV area increased seroprevalence for raccoon populations characterized as habitat specialists or strict generalists, but not for raccoons displaying weak habitat selection patterns similar to those exhibited by radio-tracked individuals. Raccoons are commonly described as ecological generalists, and are widespread in rural and agricultural areas, prairie ecosystems, and urban and suburban landscapes (Prange et al. 2003, Chamberlain et al. 2007, Beasley et al. 2011). At the population scale, our resource selection analysis suggested weak resource selection among raccoons for wooded areas and wetlands, consistent with other resource selection analyses in raccoons conducted across a variety of landscapes (Urban 1970, Prange et al. 2003, Chamberlain et al. 2007). Yet we also report substantial individual-level variation in habitat selection strength, as well as a wide range of individual responses to a variety of habitats among radio-tracked raccoons. This variation may reflect behavioral plasticity in response to fine-scale environmental variability in resources across urban-suburban landscapes or could reflect heterogeneity among raccoons in their degree of individual specialization (Piersma and Drent 2003, Schuttler et al. 2015). Individual-level variation in habitat selection could reduce achieved seroprevalence using habitat-based baiting designs by dampening the effects of a population-level baiting design that uses population averages like the one we modeled. Our results underscore the utility of leveraging variable individual habitat selection and avoidance to identify different behavioral tactics and their intrinsic and extrinsic drivers (Bastille-Rousseau and Wittemyer 2019), which could then be used to refine baiting to target specific combinations of strategies.

Our results suggest several promising avenues for optimizing the spatial distribution of baits targeting meso-carnivores in urban-suburban landscapes. First, we found that seroprevalence gains were highest with area-wide baiting relative to road-limited baiting across all baiting designs all else being equal, because more of the raccoon-preferred habitats were away from roads. In rural and agricultural areas, the standard approach to bait delivery is by fixed-wing aircraft employing GPS technologies and automated machinery to standardize and accurately deliver baits along pre-programmed and precisely spaced flight lines (Slate et al. 2009). In densely populated urban-suburban landscapes, however, fixed-wing aircraft delivery is not feasible, and NRMP baiting is accomplished predominantly by hand-baiting from vehicles along roadways (Slate et al. 2009) Our work indicates that alternative delivery methods like helicopters or bait stations which extend bait application beyond major roads in developed areas could improve baiting effectiveness and vaccination coverage (Berentsen et al. 2018, Slate et al. 2020). In addition, spatial bait coverage along roads could be further improved by incorporating minor roads into delivery planning in addition to major roads. This approach is currently being implemented by NRMP by using smaller subunits for baiting targets (i.e., planning using 1 km^2^ rather than 37 km^2^ subunits). However, an important implication of our artificial landscape simulation results is that habitat-based baiting can increase seroprevalence along roads dramatically, when the roads pass through heterogeneous and highly available raccoon-preferred habitats.

A second point for spatial bait optimization is that bait density played an important role in predicting seroprevalence in our models, as expected. Rabies seroprevalence decreased significantly when bait density was halved, leading to a reduction of seroprevalence from 35% to 24% on average. The current target density for bait application in rural and urban-suburban ORV zones in the U.S. and Canada ranges from 37.5 - 300 baits/km^2^, with 75 (rural) and 150 baits/km^2^ (urban-suburban) being standard application rates (Slate et al. 2020). Our work suggests that the current target bait density of 150 baits/km^2^ may be the minimum lower threshold needed for effective baiting in urban areas with moderate to high raccoon densities (5-30 raccoons/km^2^). Given that baits are the most expensive cost associated with ORV baiting, the optimization of bait density represents an important avenue for increasing both cost and population-level vaccination coverage effectiveness (Slate et al. 2020). Third, we found that increasing the spatial coverage of baits from 3% to 6% within the baiting area led to notable increases in seroprevalence, suggesting that even within existing road delivery methods, baiting modifications that increase coverage (i.e., evenness of bait distribution) could yield significant gains in baiting effectiveness, provided they are logistically feasible. Finally, we note that although alternative baiting strategies increased predicted seroprevalence in our models relative to standard NRMP conditions, maximum predicted seroprevalence remained below 50%. These predicted seroprevalence levels are lower than the minimum predicted vaccination coverage (seroprevalence) needed to induce population-level herd immunity based on rabies modeling efforts (Thulke and Eisinger 2008, Rees et al. 2013, Reynolds et al. 2015). We note that a recent evaluation of RRV elimination status that included the Burlington, Vermont area suggests ORV- associated case reduction in the context of declining regional prevalence (Davis et al. 2019). There may be additional spatial factors at play regarding the landscape transition from rural to urban areas which impact RV control in raccoon populations in the northeastern U.S., in part to explain why apparently suboptimal seroconversion may still be associated with case reduction in urban areas.

Urban development and land use change, including roads, influence landscape ecology and may impact target animal movements, space use, and bait availability. Roads can act as barriers to or facilitate movement in wildlife, either increasing habitat fragmentation or increasing landscape connectivity, depending on species and population-specific behavioral responses (Coffin 2007). Raccoons may avoid or be averse to crossing major roads while heavily utilizing culverts and other road-associated anthropogenic habitats for denning and foraging sites along smaller roads (Hoffmann and Gottschang 1977, Prange et al. 2003). Major roads could constrain raccoon movement between large patches of preferred habitat through avoidance behaviors, decreasing the accessibility of habitats to raccoons and diminishing the effectiveness of habitat-based baiting where baits are delivered on roads. Alternatively, raccoons may utilize road-associated habitat indiscriminately, with the effects of roads on baiting effectiveness driven by road and landscape composition and their interactions with raccoon movement and habitat selection. We did not include raccoon behavioral responses to roads in our models, but our habitat selection analysis uncovered significant and consistent avoidance behaviors of raccoons to medium and high intensity development, which correlates with major road structure. An explicit consideration of how urban-adapted meso-carnivores respond to roads, with a specific emphasis on whether roads influence movements and habitat use in a manner that enhances or decreases bait encounter rates, could provide insight into optimal bait application along roads. Finally, given that gains in seroprevalence from habitat-based baiting were greater in artificial landscapes for data-based raccoon movement compared to the actual Burlington landscape, our artificial landscape analysis highlights the limits of generalizing results to other landscapes without a better understanding of how habitat selection among target wildlife interacts with road and landscape structure

Spatial heterogeneity is both a cause and consequence of physical and biological processes that could influence wildlife disease management efforts using oral baiting strategies. Landscape heterogeneity can directly affect the movement and spatial distribution of wildlife at both broad and fine spatial scales—including for raccoons and other urban-adapted meso-carnivores like striped skunks and coyotes (Russell et al. 2005, Cullingham et al. 2009, Rioux-Paquette et al. 2014, Tardy et al. 2014). Spatial heterogeneities can also emerge from dynamic ecological processes such as sociality and demography. In our models, we included density-dependent conspecific avoidance because it can be an important cue for individual-level movement decisions and because conspecific density can mediate space use through density-dependent effects on habitat selection, including in raccoons (Tardy et al. 2014, White et al. 2018). However, we did not include social or family group behaviors in our models. Spatial heterogeneity in host-host contacts stemming from social behaviors could affect conspecific bait competition in a dynamic and non-linear way that we did not capture in this work. Furthermore, contact heterogeneity can also have important implications for a directly transmitted pathogen like RV (Reynolds et al. 2015). Raccoons exhibit a highly variable fission-fusion social structure that varies with seasonality, habitat availability and provisioning, and conspecific density (Gehrt and Fritzell 1998, Cullingham et al. 2008, Dharmarajan et al. 2009). Conspecific behavior and social structure in raccoons may differ in urban environments relative to rural areas. Additional extensions of our modeling framework could thus include social structure and conspecific behavior in urban-adapted raccoons. An additional key extension to our modeling framework would be the inclusion of RV transmission and disease spread to explore the mechanistic effects of habitat selection and landscape heterogeneity on disease incidence and population-level ORV effectiveness.

Our approach leverages methods in movement ecology to improve zoonotic disease management efforts and incorporates raccoon-urban habitat associations through movement and habitat selection analyses informed by raccoon movement data. We implemented a 3^rd^ order within-home range RSF for modeling raccoon movement that was informative at the spatial scale that baiting is conducted and could be flexibly applied to other wildlife and landscapes. RSFs provide an objective relative ranking of habitat or resources compared to ranking based on expert opinion often used in wildlife baiting programs. However, we note that depending on the species and movement parameters of interest, other approaches might also be applicable. For example, integrated Step Selection Analyses (iSSAs, Avgar et al. 2016) could be an efficient way of parameterizing movement-based IBMs given iSSAs estimate movement parameters and selection coefficients in a single step, making the coefficients potentially more robust and informative (but see also Bastille-Rousseau et al. 2018 on potential limits of such an analysis in informing movement behavior). In addition, the conditional nature of the iSSA means that the coefficients representing selection reflect how landcovers are being selected locally. However, the benefits of an integrated approach such as an iSSA might be diminished if other key ecological processes are modeled within the IBM framework, such as in our case with home range center attraction and avoidance of conspecifics, as these two aspects are not directly evaluated in an iSSA approach. Nevertheless, a robust assessment of alternative approaches to modeling movement and wildlife-habitat associations that could enhance efficacy of oral vaccination campaigns and other wildlife baiting programs would be of merit and valuable in future work.

Spatially explicit models can be useful for testing alternative control and mitigation strategies within an adaptive management paradigm in which infectious disease management and research are iterative (White and Forester 2018, Gilbert and Chipman 2020). Our work supports the idea that a non-uniform landscape-level application of baits can be effective, as is reflected in current expert opinion-based NRMP baiting strategies, and as suggested by others (Eisinger et al. 2005, Russell et al. 2005, Haydon et al. 2006, Boyer et al. 2011) We also show that the specific benefits of habitat baiting on seroprevalence may be context specific, with the degree and variability in habitat selection interacting with urban-suburban landscape composition and patch size to determine population seroprevalence. Our work thus underscores the utility of accounting for both habitat selection and landscape composition and structure in planning baiting efforts, and supports the continued prioritization of movement data collection within ORV programs targeting wildlife to better understand habitat selection and movement patterns in urban-suburban landscapes, especially during baiting season(s). Our novel simulation approach could readily be applied to other baiting and target species systems to serve as a tool for bait optimization, in support of effective oral baiting for invasive species and wildlife disease management.

## ACKNOWLEDGEMENTS

We thank the USDA National Rabies Management Program for supporting this work and for fruitful modeling discussions with the USDA National Wildlife Research Center’s quantitative group. K.M.M., K.M.P., and A.T.G. designed the research with input from G.B.R., Z.A., G.W.; K.M.M. and G.B.R. wrote the model code with contributions from A.J.D. and K.M.P.; G.B.R. analyzed the movement data, conducted the simulations, and analyzed the model results with input from K.M.P; C.A.S. collected raccoon telemetry data; K.N. and R.B.C. led ORV efforts and bait data collection; K.M.M., K.M.P., and G.B.R. wrote the first draft of the manuscript; all authors provided substantial critical feedback on the draft and gave final approval for publication. The findings and conclusions in this publication have not been formally disseminated by the USDA and do not represent Agency determination or policy.

## DATA AVAILABILITY

Model code is available via the Dryad Digital Repository http://dx.doi.org/…

## Appendix 1

### Supplementary Information

Appendix 1 includes

Figures S1 – S3

Tables S1 – S3

**Table S1.**
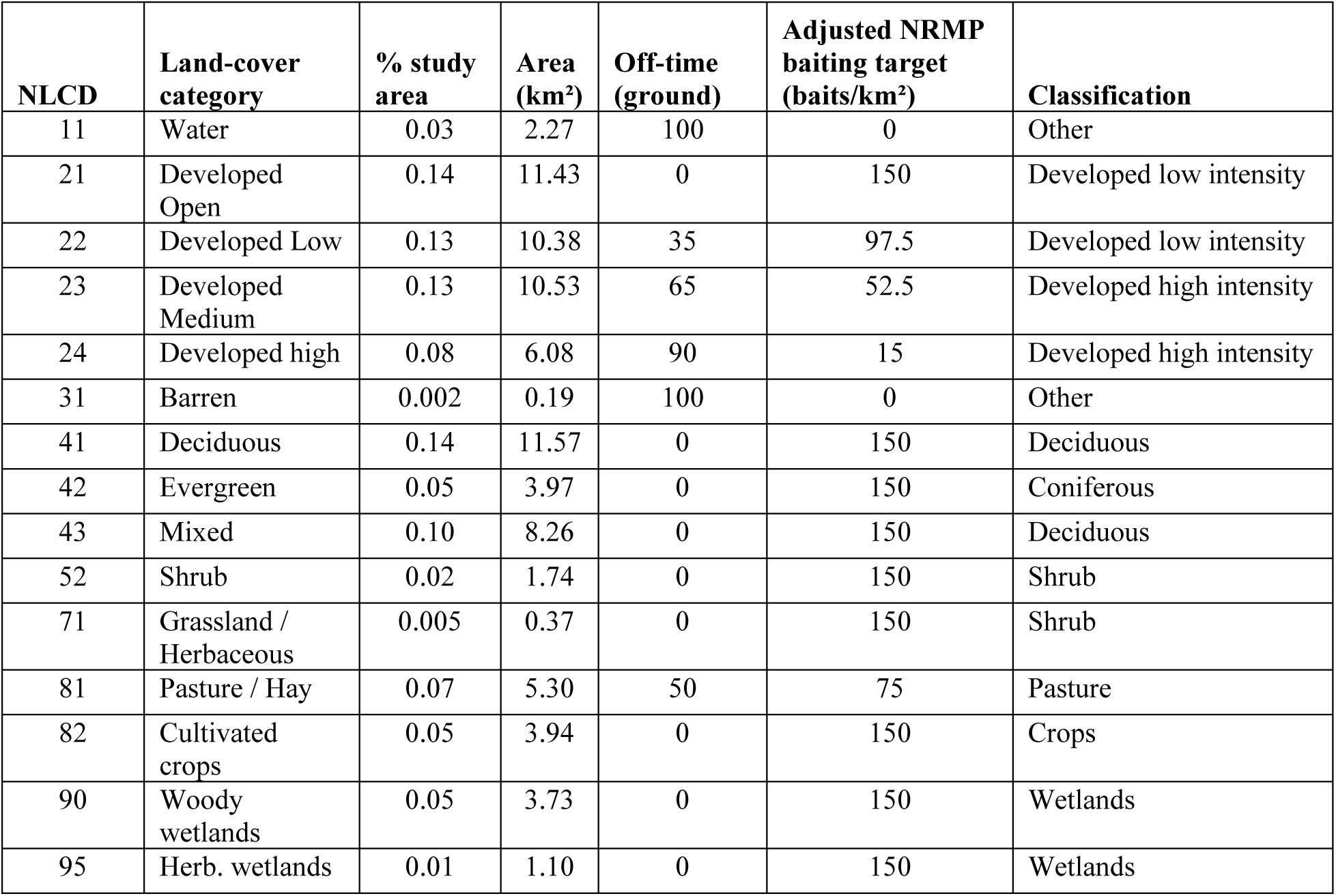
Baiting delivery algorithm. Landscape composition, National Rabies Management Program (NRMP) baiting algorithms, and landscape classification used in habitat selection analysis for the 81km^2^ Burlington, Vermont, USA study area. NLCD refers to the Multi-resolution Land Characteristics Consortium’s National Land Cover Data 2011 classifications. Off-time is the percentage of time during which bait delivery is restricted in this land cover type.

**Table S2.** Habitat selection analysis results. Individual-level resource selection coefficients and step length distribution parameters for each raccoon and land cover type in the greater Burlington, Vermont area. RSF coefficients are relative to low-intensity urban development as classified using NLCD land cover data. Shape and scale are parameter estimates of the gamma distribution fit to individual-level step length distances (in m) at a 30-minute interval. Population-level coefficients for data-based (Pop.) and theoretical habitat specialist (Specialist) are also shown.

| ID | Shape | Scale | Low Dev. | Urban | Deciduous | Coniferous | Shrub | Pasture | Crops | Wetland | Water |
| --- | --- | --- | --- | --- | --- | --- | --- | --- | --- | --- | --- |
| 1 | 0.81 | 160.40 | 0 | -1.466 | 0.675 | 0 | 1.269 | 1.854 | 0.412 | 0.635 | -10.899 |
| 3 | 0.97 | 151.70 | 0 | -0.106 | -0.439 | -15.827 | -15.827 | 0 | 0 | 1.793 | -15.827 |
| 4 | 0.71 | 215.28 | 0 | -2.324 | -0.736 | -0.570 | 0.089 | -1.057 | -2.081 | 0.690 | -16.214 |
| 5 | 1.09 | 83.06 | 0 | -0.684 | -0.076 | -0.992 | -1.231 | 0 | 0 | -1.268 | -16.905 |
| 6 | 0.94 | 59.02 | 0 | -0.567 | 0 | 0 | 0 | 0 | 0 | -0.469 | -16.207 |
| 7 | 1.33 | 80.87 | 0 | -0.957 | 0.421 | 14.027 | 1.448 | 0 | 0 | -0.078 | 0 |
| 8 | 1.02 | 143.96 | 0 | -0.266 | 0.489 | 0 | -0.840 | 0 | 0 | 2.241 | -15.879 |
| 9 | 0.79 | 64.71 | 0 | -0.255 | 20.291 | 0 | -14.841 | 0 | 0 | 0 | -14.841 |
| 10 | 1.13 | 80.62 | 0 | -0.984 | 0.781 | 0.207 | -15.766 | 0 | 0 | -0.973 | -15.766 |
| 11 | 0.86 | 80.15 | 0 | -0.355 | 0 | 0 | -2.032 | 0 | 0 | 0 | -15.978 |
| 12 | 0.99 | 66.13 | 0 | 0.004 | 0 | 0 | 0 | 0 | 0 | 0 | 0 |
| 13 | 0.96 | 68.26 | 0 | 0.008 | 0 | 0 | 0 | 0 | 0 | 0 | 0 |
| 14 | 0.67 | 125.21 | 0 | -0.209 | 0 | 0 | 0 | 0 | 0 | 0 | -15.116 |
| 15 | 0.74 | 195.13 | 0 | -0.769 | 0.614 | -0.707 | 2.882 | -14.771 | -14.771 | 0.550 | 0 |
| 16 | 0.87 | 205.59 | 0 | -1.688 | -0.076 | 0.447 | 0.149 | -0.767 | -1.385 | 0.561 | 0 |
| 17 | 0.72 | 111.60 | 0 | 1.062 | 0.155 | 0.910 | 0.956 | -1.626 | 0 | 1.122 | -13.413 |
| 19 | 0.89 | 210.51 | 0 | -1.314 | -0.626 | -0.172 | -0.269 | -0.798 | -0.490 | -0.366 | 0 |
| 20 | 0.85 | 95.52 | 0 | -0.366 | 0.139 | -1.433 | -0.023 | 0 | 0 | -0.125 | -14.119 |
| 21 | 0.95 | 178.30 | 0 | -1.042 | 0.281 | 0.399 | 0.666 | -0.653 | -0.034 | -0.043 | 0 |
| 22 | 0.66 | 237.34 | 0 | -0.421 | -0.017 | -14.775 | 1.143 | -1.559 | -14.775 | 0.138 | -14.775 |
| 23 | 1.02 | 92.02 | 0 | -13.383 | 0.437 | -0.912 | -1.119 | -2.193 | 0.349 | 1.330 | 0 |
| 24 | 0.66 | 201.10 | 0 | -0.770 | 0.385 | -0.187 | -0.680 | -0.512 | 20.945 | -0.071 | -16.188 |
| 26 | 1.37 | 151.59 | 0 | -2.124 | -0.002 | -0.281 | -0.739 | -14.890 | 0 | 0.954 | 0 |
| 28 | 0.99 | 174.28 | 0 | -1.549 | 0.232 | 0.236 | -0.112 | -0.972 | -0.314 | 1.106 | 0 |
| 29 | 0.85 | 162.58 | 0 | -1.324 | -0.756 | -13.496 | -1.941 | -1.744 | 0 | -0.936 | -3.753 |
| **Pop.** | 0.78 | 151.98 | 0 | -0.496 | 0.124 | -0.143 | 0.441 | -0.044 | -0.556 | 0.56 | -6.464 |
| **Specialist** | 0.78 | 151.98 | 0 | 0 | 0.817 | 0 | 0 | 0 | 0 | 1.253 | 0 |

**Table S3.** Rabies seroprevalence predictions. Effects of baiting design and raccoon habitat selection on predicted rabies seroprevalence as estimated using a mixed-effects logit regression. Coefficients indicate whether covariates (or their interactions) increase or decrease seroprevalence. Each covariate was categorical with the reference level (the intercept) set as raccoon movement based on individual data (GPSInd), bait distribution based on actual data (BaitDataBased), 110 baits/km^2^ (Bait110), 15 raccoons/km^2^ (Raccoon15), baits limited to road (RoadT), and with 3% spatial coverage (Cvg3). LCI and UCI correspond to lower and upper confidence intervals, respectively. Marginal and conditional R^2^ shown.

| Model | Coefficient | LCI | UCI |
| --- | --- | --- | --- |
| Intercept (GPSInd, BaitDataBased, Raccoon15, Bait110, RoadT, Cvg3) | -0.63309 | -0.70073 | -0.56546 |
| Random | 0.214501 | 0.118849 | 0.310153 |
| GPSPop | 0.18507 | 0.089418 | 0.280722 |
| SpecialistPop | -0.34103 | -0.43669 | -0.24538 |
| SpecialistInd | -0.44249 | -0.53814 | -0.34683 |
| BaitRandom | 0.06232 | -0.03333 | 0.157972 |
| BaitRSF (Specialist) | 0.348962 | 0.25331 | 0.444614 |
| BaitRSF (Data based) | -0.10461 | -0.20026 | -0.00896 |
| Raccoon30 | -0.1927 | -0.26034 | -0.12506 |
| Raccoon5 | 0.173329 | 0.105693 | 0.240966 |
| Bait55 | -0.4951 | -0.55032 | -0.43987 |
| RoadF | 0.618933 | 0.563709 | 0.674158 |
| Cvg6 | 0.186625 | 0.1314 | 0.24185 |
| Random:BaitRandom | 0.069873 | -0.0654 | 0.205146 |
| GPSPop:BaitRandom | 0.044967 | -0.09031 | 0.18024 |
| SpecialistPop:BaitRandom | -0.0481 | -0.18337 | 0.087174 |
| SpecialistInd:BaitRandom | -0.01979 | -0.15506 | 0.115483 |
| SpecialistPop:BaitRSF | -0.0715 | -0.20678 | 0.06377 |
| GPSPop:BaitRSF | -0.05926 | -0.19453 | 0.076012 |
| GPSInd:BaitDataBased:Raccoon30 | 0.022375 | -0.07328 | 0.118027 |
| Random:BaitDataBased:Raccoon30 | 0.026374 | -0.06928 | 0.122026 |
| GPSPop:BaitDataBased:Raccoon30 | 0.007987 | -0.08767 | 0.103639 |
| SpecialistPop:BaitDataBased:Raccoon30 | 0.017603 | -0.07805 | 0.113255 |
| SpecialistInd:BaitDataBased:Raccoon30 | 0.019965 | -0.07569 | 0.115617 |
| GPSInd:BaitRandom:Raccoon30 | 0.032415 | -0.06324 | 0.128067 |
| Random:BaitRandom:Raccoon30 | 0.028927 | -0.06673 | 0.124579 |
| GPSPop:BaitRandom:Raccoon30 | 0.008786 | -0.08687 | 0.104439 |
| SpecialistPop:BaitRandom:Raccoon30 | 0.035104 | -0.06055 | 0.130757 |
| SpecialistInd:BaitRandom:Raccoon30 | 0.0285 | -0.06715 | 0.124152 |
| SpecialistPop:BaitRSF:Raccoon30 | -0.00745 | -0.1031 | 0.088203 |
| SpecialistInd:BaitRSF:Raccoon30 | -0.00904 | -0.10469 | 0.086612 |
| GPSInd:BaitRSF:Raccoon30 | 0.022433 | -0.07322 | 0.118085 |
| GPSInd:BaitDataBased:Raccoon5 | -0.03465 | -0.13031 | 0.060998 |
| Random:BaitDataBased:Raccoon5 | -0.04495 | -0.1406 | 0.050705 |
| GPSPop:BaitDataBased:Raccoon5 | -0.01437 | -0.11002 | 0.081287 |
| SpecialistPop:BaitDataBased:Raccoon5 | -0.01278 | -0.10843 | 0.082876 |
| SpecialistInd:BaitDataBased:Raccoon5 | -0.0107 | -0.10636 | 0.084949 |
| GPSInd:BaitRandom:Raccoon5 | -0.0325 | -0.12815 | 0.063152 |
| Random:BaitRandom:Raccoon5 | -0.03955 | -0.1352 | 0.056107 |
| GPSPop:BaitRandom:Raccoon5 | -0.01942 | -0.11507 | 0.076231 |
| SpecialistPop:BaitRandom:Raccoon5 | -0.01107 | -0.10672 | 0.084583 |
| SpecialistInd:BaitRandom:Raccoon5 | -0.00137 | -0.09703 | 0.094279 |
| SpecialistPop:BaitRSF:Raccoon5 | 0.008979 | -0.08667 | 0.104631 |
| SpecialistInd:BaitRSF:Raccoon5 | 0.01225 | -0.0834 | 0.107902 |
| GPSInd:BaitRSF:Raccoon5 | -0.02982 | -0.12547 | 0.065832 |
| GPSInd:BaitDataBased:Bait55 | -0.03389 | -0.11199 | 0.044211 |
| Random:BaitDataBased:Bait55 | -0.0492 | -0.1273 | 0.028902 |
| GPSPop:BaitDataBased:Bait55 | -0.05412 | -0.13222 | 0.023977 |
| SpecialistPop:BaitDataBased:Bait55 | -0.04356 | -0.12166 | 0.03454 |
| SpecialistInd:BaitDataBased:Bait55 | -0.02755 | -0.10565 | 0.050552 |
| GPSInd:BaitRandom:Bait55 | -0.04015 | -0.11825 | 0.037953 |
| Random:BaitRandom:Bait55 | -0.05673 | -0.13483 | 0.021368 |
| GPSPop:BaitRandom:Bait55 | -0.06025 | -0.13835 | 0.01785 |
| SpecialistPop:BaitRandom:Bait55 | -0.04123 | -0.11933 | 0.036873 |
| SpecialistInd:BaitRandom:Bait55 | -0.03082 | -0.10892 | 0.047279 |
| SpecialistPop:BaitRSF:Bait55 | 0.006045 | -0.07205 | 0.084145 |
| SpecialistInd:BaitRSF:Bait55 | 0.022992 | -0.05511 | 0.101092 |
| GPSInd:BaitRSF:Bait55 | 0.02233 | -0.05577 | 0.10043 |
| GPSInd:BaitDataBased:RoadF | -0.06918 | -0.14728 | 0.008919 |
| Random:BaitDataBased:RoadF | -0.36931 | -0.44741 | -0.29121 |
| GPSPop:BaitDataBased:RoadF | -0.2791 | -0.3572 | -0.201 |
| SpecialistPop:BaitDataBased:RoadF | 0.320047 | 0.241947 | 0.398146 |
| SpecialistInd:BaitDataBased:RoadF | 0.399975 | 0.321875 | 0.478074 |
| GPSInd:BaitRandom:RoadF | -0.19311 | -0.27121 | -0.11501 |
| Random:BaitRandom:RoadF | -0.41957 | -0.49767 | -0.34147 |
| GPSPop:BaitRandom:RoadF | -0.41215 | -0.49025 | -0.33405 |
| SpecialistPop:BaitRandom:RoadF | 0.135141 | 0.057042 | 0.213241 |
| SpecialistInd:BaitRandom:RoadF | 0.187631 | 0.109531 | 0.265731 |
| SpecialistPop:BaitRSF:RoadF | 0.165767 | 0.087667 | 0.243866 |
| SpecialistInd:BaitRSF:RoadF | 0.191732 | 0.113632 | 0.269832 |
| GPSInd:BaitRSF:RoadF | 0.074 | -0.0041 | 0.1521 |
| GPSInd:BaitDataBased:Cvg6 | 0.171003 | 0.092903 | 0.249102 |
| Random:BaitDataBased:Cvg6 | 0.191254 | 0.113155 | 0.269354 |
| GPSPop:BaitDataBased:Cvg6 | 0.185023 | 0.106923 | 0.263123 |
| SpecialistPop:BaitDataBased:Cvg6 | 0.114444 | 0.036344 | 0.192544 |
| SpecialistInd:BaitDataBased:Cvg6 | 0.110162 | 0.032062 | 0.188262 |
| GPSInd:BaitRandom:Cvg6 | 0.046018 | -0.03208 | 0.124117 |
| Random:BaitRandom:Cvg6 | 0.053673 | -0.02443 | 0.131773 |
| GPSPop:BaitRandom:Cvg6 | 0.056382 | -0.02172 | 0.134482 |
| SpecialistPop:BaitRandom:Cvg6 | 0.04555 | -0.03255 | 0.123649 |
| SpecialistInd:BaitRandom:Cvg6 | 0.045059 | -0.03304 | 0.123159 |
| SpecialistPop:BaitRSF:Cvg6 | -0.00771 | -0.08581 | 0.070389 |
| SpecialistInd:BaitRSF:Cvg6 | -0.01323 | -0.09133 | 0.064869 |
| GPSInd:BaitRSF:Cvg6 | 0.00027 | -0.07783 | 0.07837 |
| **Marginal R<sup>2</sup>** | **0.956314** |  |  |
| **Conditional R<sup>2</sup>** | **0.982438** |  |  |

**Figure S1.**
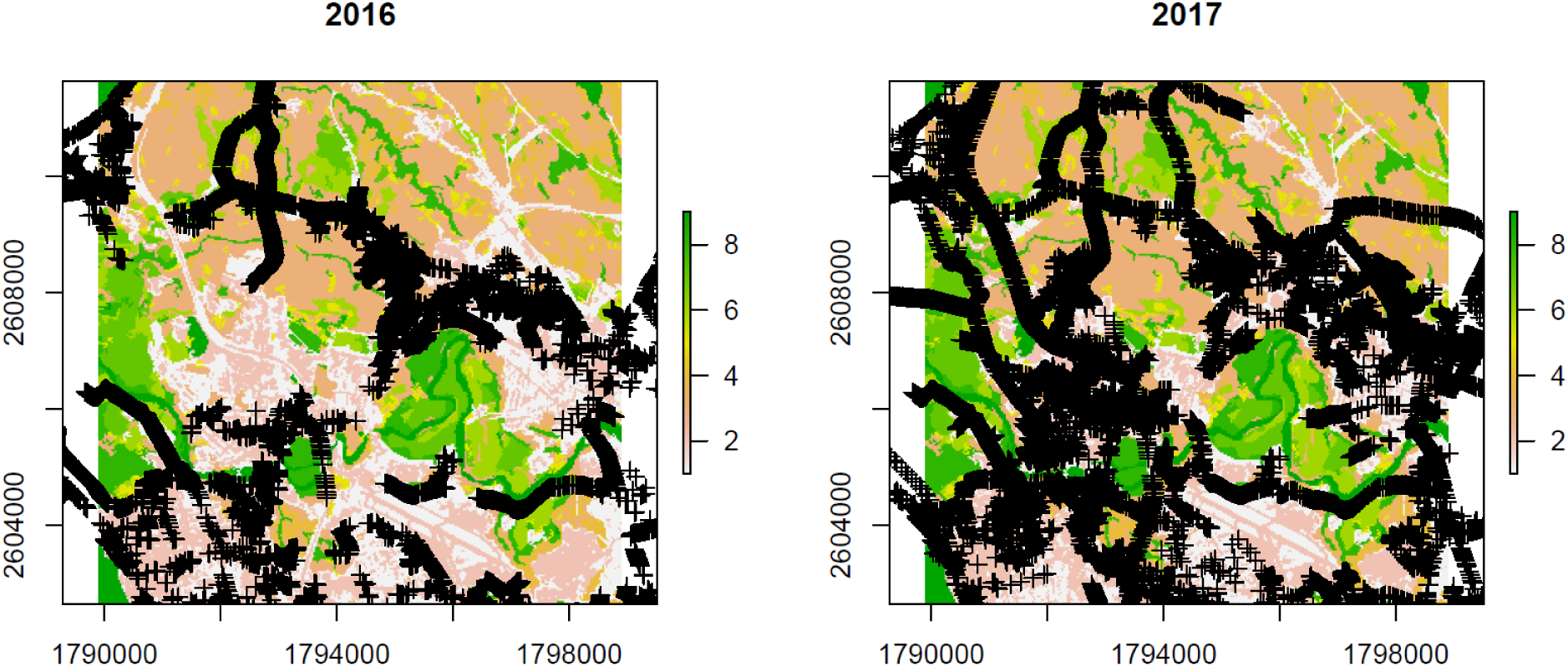
ORV bait distribution in Burlington, Vermont. GPS bait locations deployed by NRMP in 2016 and 2017 along roads are shown in black. Land cover reclassification used in the habitat selection analysis is indicated by the color ramp. Bait density was 150 baits/km^2^ in target landcover classes both years. Grid cell coverage at the 900 m^2^ resolution used in simulations was 3% in 2016 to 6% in 2017.

**Figure S2.**
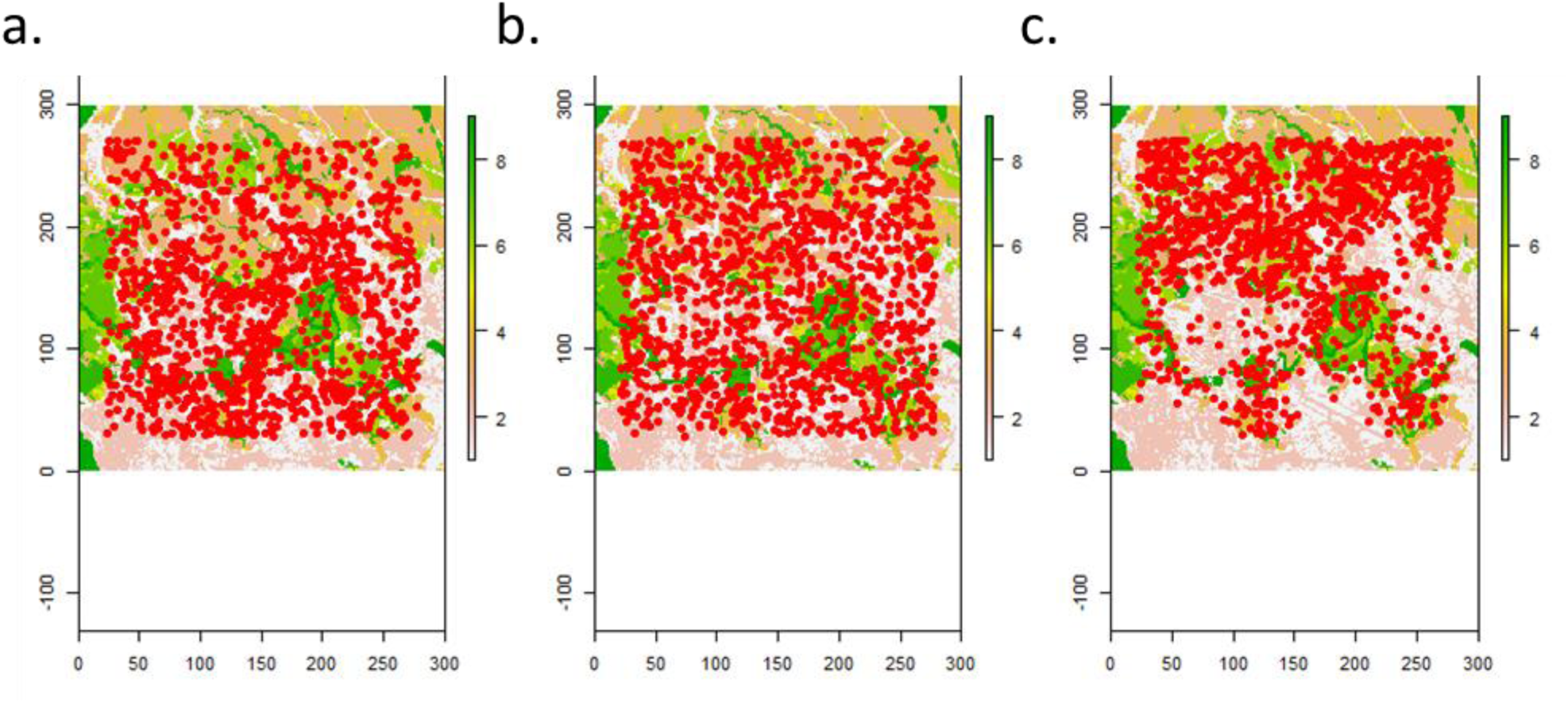
Initialization of raccoons in movement simulations. Red dots represent raccoons prior to the start of the movement simulations. Examples of initialization of individuals shown for: a) data-based habitat selection; b) strict generalist, with starting locations for individuals generated at random; and c) specialist, with starting locations biased towards wetland and deciduous forest.

**Figure S3.**
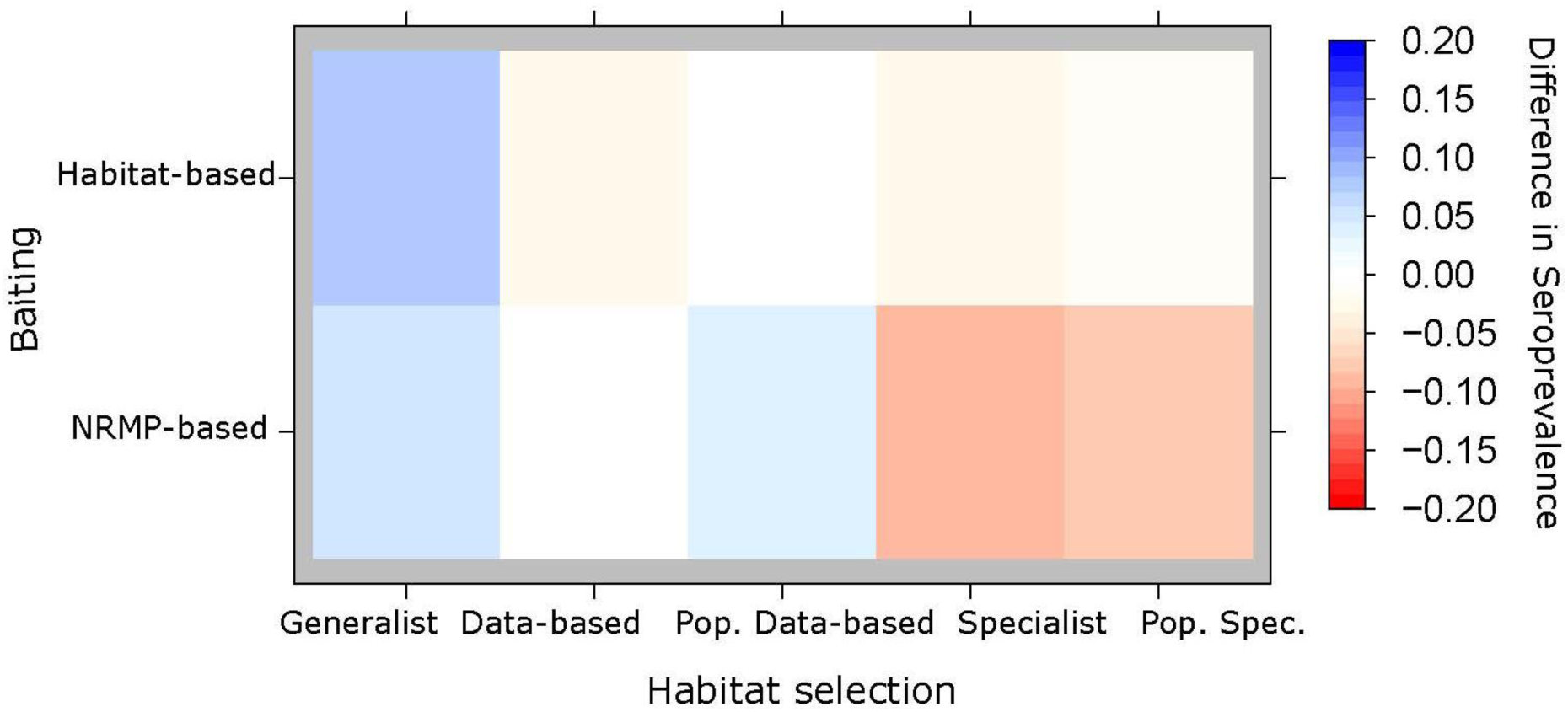
Effects of individual-level variation in habitat selection on seroprevalence. Plots show absolute difference in seroprevalence for different baiting designs relative to National Rabies Management Program (NRMP) baiting conditions (white square located at [2,2] in the panel; predicted absolute seroprevalence of 0.35). Rows in each panel correspond to the spatial distribution of baits across land cover types (i.e. proportion of baits distributed to different landcover types). Habitat-based baiting stratifies baiting using habitat selection patterns while NRMP-based baiting stratifies baiting using an algorithm drawn from expert opinion by the oral rabies vaccination team at NRMP. Columns correspond to raccoon habitat selection behaviors, including individual-level variation in habitat selection (Generalist, Data-based, and Specialist) and population-level habitat selection (Pop. Data-based and Pop. Specialist).

